# A Critical Study in Stereopsis and Listing’s Law

**DOI:** 10.1101/2023.07.04.547695

**Authors:** Jacek Turski

## Abstract

The brain uses slightly different 2D retinal images to enhance our vision with stereopsis: spatial depth and 3D shape. Stereopsis is organized by pairs of corresponding retinal elements of zero disparity: a small retinal area in one eye and the corresponding unique area in the other share one subjective visual direction. This organization results in retinal disparity’s spatial coordinates. The study presented here extends the 2D setting of the author’s geometric modeling of the disparity’s spatial coordination in the binocular system with the asymmetric eye (AE) to the 3D framework. The AE models the healthy human eye’s asymmetry of optical components. In *GeoGebra’s* dynamic geometry simulations, the 3D spatial coordinates of retinal disparity integrated with the eyes’ posture are visualized, and longitudinal and vertical disparities of distal visual stimuli are computed, contributing to stereopsis and visual space geometry study. Further, the torsional disparity is computed in the framework of Euler’s rotation theorem. It can assess the geometric and neural or ocular motor plant constraints to Listing’s law. Finally, epipolar geometry in the binocular system with AEs is discussed. Although this study enhances the geometric description of stereopsis and oculomotor control of eyes 3D orientations, it also simplifies their analyses.

## 1 Introduction

Slightly different 2D images of light beams reflected off the objects in space impinge on the retinal photoreceptors of our two laterally separated eyes. The brain processes these disparate stimuli to enhance our visual experience with texture, color, depth, form, and motion. However, this enhancement is task-dependent and influenced by contextual factors, learning, and expectations.

Remarkably, the binocular disparity of retinal images, as revealed by Julesz’s random-dot-stereograms, provides the single cue for our 3D visual experience, or stereopsis, with no explicit recognition of a scene’s geometric forms [Julesz, 1960, Julesz, 1964]. This realization laid the ground for studies on the physiology basis of stereopsis, cf. [Barlow et al., 1967, Hubel and Wiesel, 1962, Poggio and W.H. Talbot, 1981].

Stereopsis is organized by retinal correspondence: a small retinal area in one eye and a corresponding unique area in the other share one subjective visual direction when separately stimulated. These retinal areas are called corresponding elements determined by the nonius paradigm. We consider each pair of binocular correspondence as having zero disparity. For a given binocular fixation, the horopter is the locus of points in space such that each point on the horopter projects onto a pair of corresponding retinal elements.

The nonius measurements of the horopter revealed the asymmetry of the retinal distribution of corresponding elements: the corresponding elements are compressed in the temporal retinae relative to those in the nasal retinae [Shipley and Rawlings, 1970]. This asymmetry was one of Hillebrand’s suggestions 130 years ago to explain his measurements of the apparent frontal line, showing that the empirical horopter’s curve (although not by nonius method) deviates from the 200-year-old Vieth-Müller circular horopter (VMC) commonly referred to as geometric horopter. This discrepancy between the empirical and geometric horopters was only recently explained, as it is discussed below. The impact of this omission on the conceptual understanding of binocular vision is discussed in [Turski, 2023a].

The main tools in Julesz’s studies mentioned above were the computer-generated random-dot stereogram devised from the geometry of binocular disparity that, upon viewing it, presented us with 3D form perception. The line of inquiry that uses binocular geometry augmented by the eye’s physiological optics, integrated with the eyes’ binocular posture, and visualized in computer simulations was started in [Turski, 2018, Turski, 2020]. In those studies, the geometric theory of the longitudinal horopter closely resembling the empirical horopter was developed in the binocular system with the asymmetric eye (AE) model.

The AE model was introduced and comprehensively discussed in [Turski, 2018] and slightly modified in [Turski, 2020]. Based on the observed misalignment of optical components in healthy human eyes [Chang et al., 2007, Schaeffel, 2008], the AE comprises the fovea’s anatomical displacement from the posterior pole by angle *α* = 5.2^*°*^ and the crystalline lens’ tilt by angle *β*. Angle *α* is relatively stable in the human population, and angle *β* predominantly varies between *−*0.4^*°*^ and 4.7^*°*^. The fovea’s displacement from the posterior pole and the cornea’s asphericity contribute to optical aberrations that the lens tilt partially compensates for [Artal, 2014]. The horopter constructed by the geometrical method in [Turski, 2020] in the binocular system with AEs fully specifies the retinal correspondence asymmetry in terms of AE’s parameters *α* and *β*.

The geometric theory in [Turski, 2020] advanced the classical model of empirical horopters as conic sections introduced *ad-hoc* by Ogle in the framework of analytic geometry with free parameters for symmetric fixations [Ogle, 1932]. It was later extended by Amigo to any horizontal fixations [Amigo, 1965]. In contrast to Ogle’s theory, empirical horopters modeled in [Turski, 2020] are supported by the eye’s anatomy that contributes to the aplanatic design of the abovementioned misaligned optical components. This work was further extended to the horizontal iso-disparity (or disparity for short) conic sections, where zero-disparity conic is the horopter, and applied to study the global aspects of phenomenal spatial relations in the framework of Riemannian geometry [Turski, 2023b]. I note that the lack of a family of iso-disparity curves that would more richly describe the structure of binocular visual space was mentioned long ago in [Arditi, 1986].

This paper extends the studies in [Turski, 2018, Turski, 2020, Turski, 2023b] from the 2D setting of the binocular system with the AE model to the fully 3D framework. Here, I visualize in *GeoGebra’s* dynamic geometry simulations the 3D longitudinal disparity curves and vertical horopter integrated with the eyes’ posture using the binocular system with the new 3D AE model augmented by the vertical asymmetry of optical components. The spatial coordination of the retinal disparities and the eyes’ binocular posture, integrated with changes in fixation, allows the computation of horizontal and vertical disparities of small visual stimuli in the binocular field and torsional disparity of the eyes’ posture computed in the framework of Euler’s rotation theorem.

The disparity curves for the upright stationary head are straight frontal lines, hyperbolas, or ellipses in the visual plane, depending on the AE parameters and the fixation point location in the binocular field. The subjective vertical horopter is a straight line inclined to the visual plane. The longitudinal, vertical, and torsional disparities are computed during simulations in all of these cases. The torsional disparity for any of the eyes’ binocular posture is calculated using Euler’s rotation theorem for each eye with the constraint of maintaining bifoveal fixations. The relation of this approach to Listing’s law and its extensions is discussed in Section 7.

The *GeoGebra* applet for experimenting with the 3D disparity conic sections’ transformations under the 3D eyes’ binocular movement is included in Supplementary Material.

## 2 Retinal Correspondence and Disparity Conic Sections

### 2.1 Horizontal Retinal Correspondence and Stereopsis

A small object located away from the horopter has nonzero disparity and, in general, is perceived as two objects. However, the brain fuses disparate images into a single percept for an object close to the horopter. The brain then uses the extracted disparity to create our sense of depth relative to the horopter. Further, the brain uses the difference in the horizontal disparity between two spatial points to sense the relative depth and, hence, our perception of objects’ shapes and their locations in 3D space [Wheatstone, 1838], that is, stereopsis and visual space geometry. Wheatstone’s stereoscope demonstrated, and Julesz’s random-dot-stereogram confirmed that stereopsis is mainly supported by horizontal correspondence.

The retinal correspondence cannot be readily measured and cannot be obtained from the only one geometrical model of empirical horopters introduced in [Ogle, 1932] 90 years ago. The main reason is the beholder’s asymmetry of retinal correspondence, which impacts the shape of empirical horopters [Shipley and Rawlings, 1970, Nelson, 1977]. However, the physiological origin of retinal asymmetry has been uncertain. This situation changed when the geometric theory of the binocular system with the AE was recently developed in [Turski, 2018, Turski, 2020], which fully specifies the retinal correspondence asymmetry in terms of the asymmetry of human eye optical components. The result is the symmetric distribution of the points in the image plane of the AE covering, under projection through the nodal point, the asymmetric retinal correspondence. This symmetric correspondence was first constructed for the unique fixation in which the image planes of the AEs and, hence, the equatorial plane of the lenses’ are coplanar; it is explained in Figure 1.

**Figure 1:**
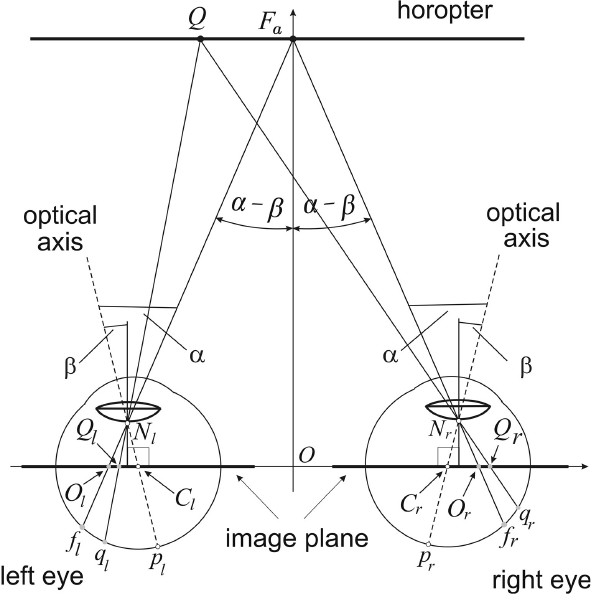
The binocular system with the AE model seen from above. For the parameters *α* and *β*, the fixation *F*_*a*_ at the abathic distance *OF*_*a*_ corresponds to the AEs posture in which the coplanar equatorial lens planes are parallel to the coplanar image planes. The point Q on the horopter projects through the nodal points *N*_*r*_ and *N*_*l*_ to the corresponding retinal points *q*_*r*_ and *q*_*l*_ and to points *Q*_*r*_ and *Q*_*l*_ in the image plane. Although |*f*_*l*_*q*_*l*_|*/*= |*f*_*r*_*q*_*r*_| on the retina, |*O*_*l*_*Q*_*l*_| = |*O*_*r*_*Q*_*r*_| on the image plane.

This unique bifoveal fixation was referred to as the eyes’ resting posture (ERP). As discussed in [Turski, 2018], the fixation in the ERP corresponds to the empirical horopter’s abathic distance fixation. Further, it numerically corresponds to the eyes’ resting vergence posture, in which the eye muscles’ natural tonus resting position serves as a zero-reference level for convergence effort [Ebenholtz, 2001].

The symmetric distribution of points on the image plane covering the asymmetric distribution of corresponding retinal points is uniquely specified by the AEs parameters *α* and *β* shown in Figure 1. The symmetric distribution is preserved for all other binocular fixations that transform the straight frontal horopter into ellipses or hyperbolas [Turski, 2020].

### 2.2 Horizontal Disparity Conic Sections

As mentioned in the previous section, the corresponding points on the image planes of the AEs are first specified in the configuration shown in Figure 1. In this figure, points *Q*_*r*_ and *Q*_*l*_ that are on the same side of the respective centers *O*_*r*_ and *O*_*l*_ and of the same distance are covering the non-uniformly distributed corresponding retinal elements *q*_*r*_ and *q*_*l*_. The corresponding points back-projected through the nodal points to physical space give the point *Q* on the horopter line.

The disparity lines in physical space are first constructed in *GeoGebra* dynamic geometry environment for the ERP; see [Turski, 2023b]. The *nδ*-disparity line is the projection of the points *Q*_*r*_ + *nδ*and *Q*_*l*_ and *− nδ*-disparity line is the set of projected points *Q*_*r*_*−nδ*and *Q*_*l*_. where *n* = 1, 2, … For *n* and *n* + 1, we have consecutive disparity objective frontal straight lines of relative disparity *δ*. The simulations of disparity lines in *GeoGebra* revealed their invariance to the abathic distance fixations and, hence, to AE’s parameters modeling asymmetry of the human eye optical components, see Figure 2 in [Turski, 2023b]. For all other binocular fixations on the points of the horizontal plane, the simulated disparity curves were either a family of ellipses or hyperbolas, depending on the AE’s parameters and the point of fixation.

**Figure 2:**
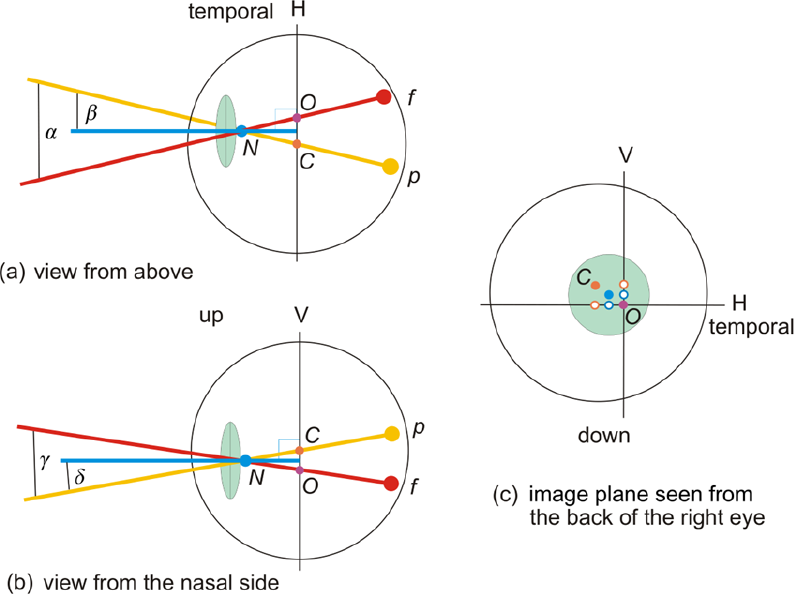
(a) The horizontal asymmetry: the fovea is displaced by angle *α* and the lens is tilted by angle *β*. (b) The vertical asymmetry: the fovea is displaced by angle *γ*, and the lens is tilted by angle *δ*. (c) The image plane is seen from the back of the right eye’s model. *H* and *V* are the horizontal and vertical coordinate axes at the image plane’s optical center *O*. The small circles on the axes in (c) show the horizontal and vertical asymmetry components. The yellow lines represent the eye’s optical axis, the red lines the visual axis, and the blue lines the lens’ optical axis. *N* is the nodal point, *C* is the eye’s center of rotation, *p* is the posterior pole, and *f* is the fovea.

## 3 The Binocular System with 3D AEs

### 3.1 3D AE model

The AE model parameters of the horizontal components of ocular asymmetry, the displacement of the fovea from the posterior pole *α* and the lens tilt *β*, are here extended by the vertical components *γ* and *ε*, respectively. The 3D AE model is shown schematically in Figure 2.

Similarly to the AE model proposed in [Turski, 2018], the 3D AE model incorporates only those elements that are indispensable in modeling the human eyes’ asymmetry of the optical components. Thus, it comprises the three axes necessary to specify the asymmetry: the eyeball’s optical axis (yellow), the visual axis (red), and the lens’ optical axis (blue).

The average anthropomorphic values of the asymmetry angles are adapted from previous studies [Aguirre, 2019, de Castro et al., 2007, Schaeffel, 2008, Wang et al., 2019]. In these clinical studies, different measuring methods (the Scheimpflug, Purkinje, and anterior segment optical coherence tomography) were used to obtain the lens tilt relative to different reference axes (corneal topographic axis, line of sight, and papillary axis). Like in any geometric description, the reference axes are optical and visual. Both these axes cannot be measured and, therefore, cannot be used clinically. For instance, there are no tools to locate the nodal point. Consequently, the prior clinical reports need extrapolations to specify the crystalline lens’ tilt.

### 3.2 Horizontal, Vertical and Torsional Disparities

Using 3D asymmetry of the optical components, *α, β, γ*, and *ε* parameters of AE, the ERP in the binocular system is defined similarly as before for the 2D eye asymmetric model shown in Figure 1. It is specified by the fixation for which the coplanar image planes are parallel to the lens equatorial planes of the stationary upright head. Including the vertical coordinate to complement the horizontal coordinate attached to the image planes’ optical centers allows for the construction of the vertical horopter and computations of vertical disparities in addition to computed before the horizontal disparities. In a strict sense, the subjective vertical horopter is defined only for fixations in the midsagittal plane. However, it should be extended by accounting for Panum’s fusional region of the empirical vertical horopter [Amigo, 1974]. These aspects are fully discussed in Section 6.

Further, the ocular torsion of each AE is computed by adding the third coordinate axis at the optical center that is perpendicular to the image plane and, hence, parallel to the lens’ optical axis. Ocular torsion around this axis is the closest to rolling about the corneal axis. In the framework of Euler’s rotation theorem and, hence, without involving Listing’s law, the difference in ocular torsions between the right AE and the left AE gives the torsional disparity for binocular eyes’ postures. This is discussed in detail in Section 7.

## 4 3D Disparity Spatial Coordination and Eyes Posture

The average anthropomorphic values of the asymmetry angles shown in Figure 2 that are used in the simulation of disparity conic sections and the subjective vertical horopter are assumed as follows: *α* = 5.2^*°*^ and *β* = 3.3^*°*^ in the horizontal direction and *γ* = 3^*°*^ and *ε* = 1^*°*^ in the vertical direction.

### 4.1 Disparity Spatial Coordinates for ERP

The fixation point *F*_*a*_ of the ERP of the stationary upright head is uniquely determined by the parameters of the binocular system with the AEs. In the simulations, I use the average anthropomorphic values of these parameters: the interocular distance of 6.5 cm and the values of asymmetry angles listed above. Then, *F*_*a*_’s coordinates in centimeters are (0, 99.55, 3.48). The simulated disparity conic sections in this eyes’ binocular posture are straight frontal lines in the visual plane containing the nodal points and the fixation *F*_*a*_.

Figure 3 shows the disparity lines for the proximal relative disparity of *δ*= 0.005 cm, cf. Section 2.2 The details of how the disparity conic sections are constructed and disparities are calculated will be discussed in Section 6. The point *P* in the binocular field is projected into the AE image planes. The right image plane projection is *P*_*r*_(*d*_*rh*_, *d*_*rv*_) and the left image plane projection is *P*_*l*_(*d*_*lh*_, *d*_*lv*_). From the simulation computed in *GeoGebra* and displayed in Figure 3 the coordinates of projections of *P* to image planes, horizontal disparity of *P* is

**Figure 3:**
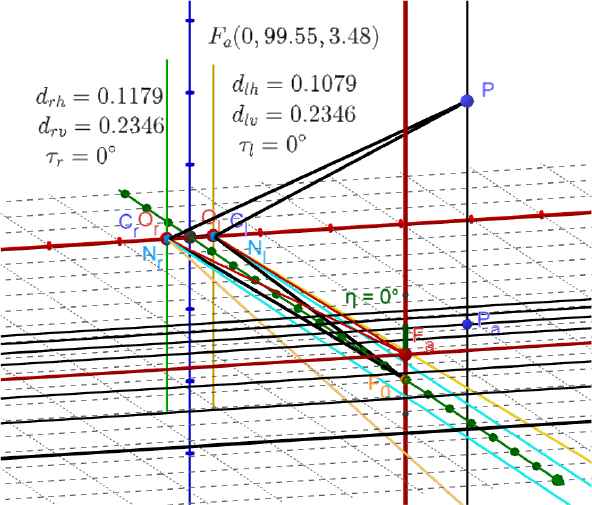
The 3D resting eyes posture with the fixation at *F*_*a*_ for the anthropomorphic AE parameters *α, β, γ*, and *δ*given in the text. The point *F*_0_ is the projection of *F*_*a*_ into the head’s transverse plane through the eyes’ rotation centers, which agrees with the horizontal plane. *N*_*r*_, *C*_*r*_, and *O*_*r*_ are the nodal point, rotation, and optical centers on the image plane, respectively, for the right AE. With the subscript “*l*,” the same points are shown for the left AE.

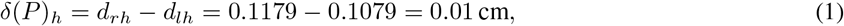

the vertical disparity is

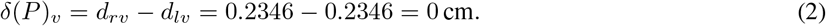

The vertical projection of point *P* on the visual plane, *P*_*a*_, is exactly on the second consecutive disparity line down from the horopter (red horizontal line through *F*_*a*_). Because the relative disparity value between each consecutive disparity line is 0.005 cm, the spatial disparity of *P* agrees with the retinal horizontal disparity *δ*(*P*)_*h*_ = 0.01 cm. The line through *P* and *P*_*a*_ is parallel to the subjective vertical horopter and agrees with the body’s anatomical axial axis indicated by *η* = 0^*°*^ but is inclined to the visual plane by about 2^*°*^.

Finally, the ocular torsions, *τ*_*r*_ and *τ*_*l*_, computed in this simulation are zero, such that the torsional disparity *δ*_*t*_ vanishes in this posture. Please refer to Section 6.2 for details on calculations of ocular torsions.

### 4.2 Disparity Spatial Coordinates for Eyes Secondary Postures

When the fixation point *F* is shifted vertically from the ERP to a secondary posture, as it is shown in Figure 4 for the down-shift from *F*_*a*_(0, 99.55, 3.48) to *F* (0, 99.55, *−*20.48), we see that the disparity lines move with the visual plane and remain straight frontal lines. Further, the subjective vertical horopter is tilted in the midsagittal plane top-away by *η* = 13.62^*°*^ at the fixation point from the objective vertical line shown by a dashed line.

**Figure 4:**
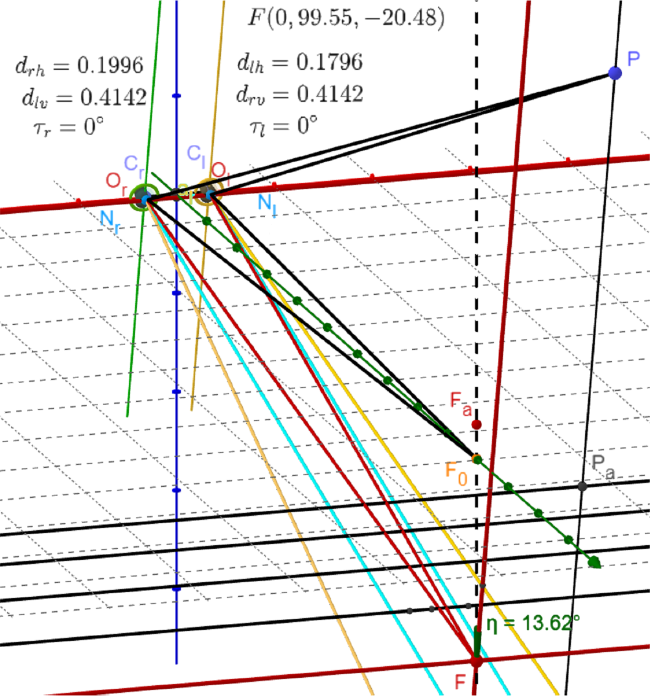
Under the vertical shift of the fixation point from the ERP, the disparity lines move with the visual plane (the plane through the nodal points and the fixation point) and remain straight frontal lines. The subjective vertical horopter is tilted from the true vertical direction. See the text for a detailed discussion.

Under this movement, the horizontal disparity of point *P* shown in this figure (this point is different than point *P* in Figure 3) changes to

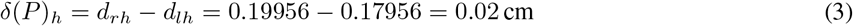

but its vertical coordinates *d*_*rv*_ and *d*_*lv*_ are the same, so *δ*(*P*)_*v*_ = 0. Further, the ocular torsions, *τ*_*r*_ and *τ*_*l*_, vanish such that the torsional disparity remains zero after the vertical shaft of *F*_*a*_.

The projection ray of *P* onto *P*_*a*_ in the visual plane is parallel to the subjective vertical horopter. It is demonstrated in Figure 4 as follows. The projection of *P* along the subjective vertical direction into the visual plane, *P*_*a*_, is on the disparity line of disparity value 4(0.005) = 0.02 cm. It agrees with the horizontal disparity value obtained from the differences of the projections in the coordinates in the right and left eye image planes and shown in (3). It demonstrates that under the objective vertical shift of the fixation point from the ERP, the visual space tilts together with the tilt of the subjective horopter by the same amount.

Further, the subjective vertical horopter tilt can be approximately calculated as follows:

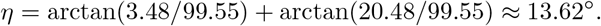

The approximate value used in the above calculations follows from the construction of the subjective vertical horopter and is explained in Section 6.1

The next secondary posture is produced by shifting *F*_*a*_(0, 99.55, 3.48) horizontally to *F* (40, 99.55, 3.48), which is shown in Figure 5.

**Figure 5:**
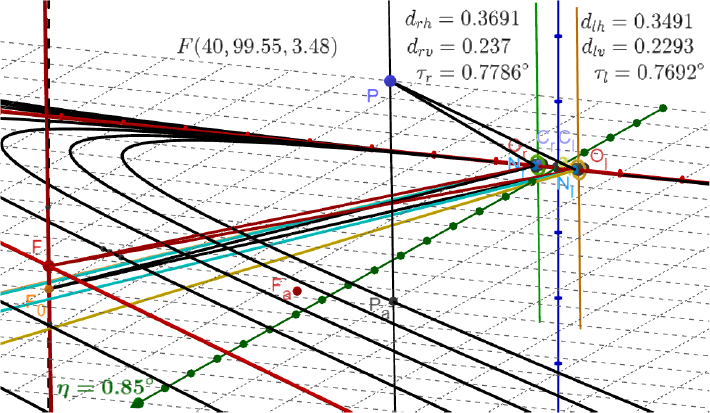
Under the horizontal shift of the fixation point from the resting eyes posture, the disparity lines in the visual plane transform into hyperbolas. The subjective vertical horopter is tilted from the true vertical direction. See the text for a detailed discussion.

Now, the disparity lines in the image plane change to hyperbolas, which is not expected for a secondary posture. Further, the subjective vertical horopter is tilted top-away by *η* = 0.85^*°*^ in the plane *−*1.7*x* + 14.7*y* = 139 containing the subjective vertical horopter (red line) and the objective vertical line (dashed line). Using the same argumentation as before, we see that the subjective vertical horopter is parallel to the line through *P* and *P*_*a*_, where *P* is chosen such that the longitudinal spatial disparity value of 4(0.005) cm agrees with retinal horizontal disparity value of *δ*_*h*_ = 0.02 cm. In contrast to the vertical secondary posture, after horizontal shift, the vertical and torsional disparities are small but not zero: *δ*_*v*_ = 0.008 cm and *δ*_*t*_ = 0.01^*°*^.

### 4.3 Disparity Spatial Coordinates for Eyes Tertiary Postures

Depending on the AE parameters and the point of fixation, the subjective disparity frontal lines, at least near the fixation, are given by families of hyperbolas or ellipses, each in the respective visual plane through the nodal and fixation points.

Figures 6 and 7 show the disparity hyperbolas and ellipses for the tertiary postures when the resting eyes fixation point *F*_*a*_ is shifted to the indicated new positions.

**Figure 6:**
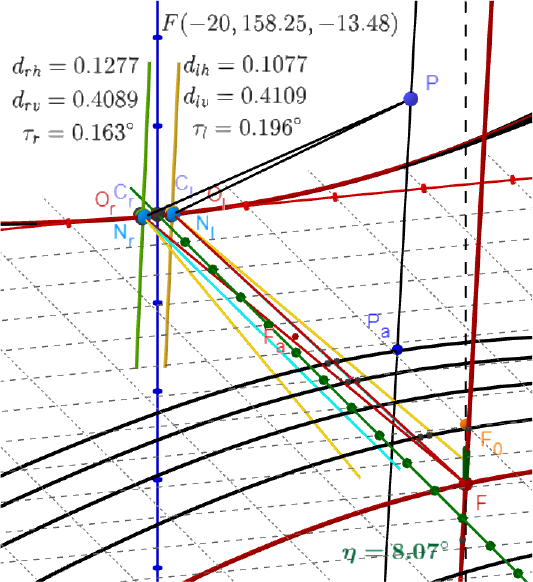
The disparity curves are hyperbolas for the fixation of the distance greater than the abathic distance. Note that the hyperbolic branches that pass through the nodal points *N*_*r*_ and *N*_*l*_ are not part of disparity curves.

**Figure 7:**
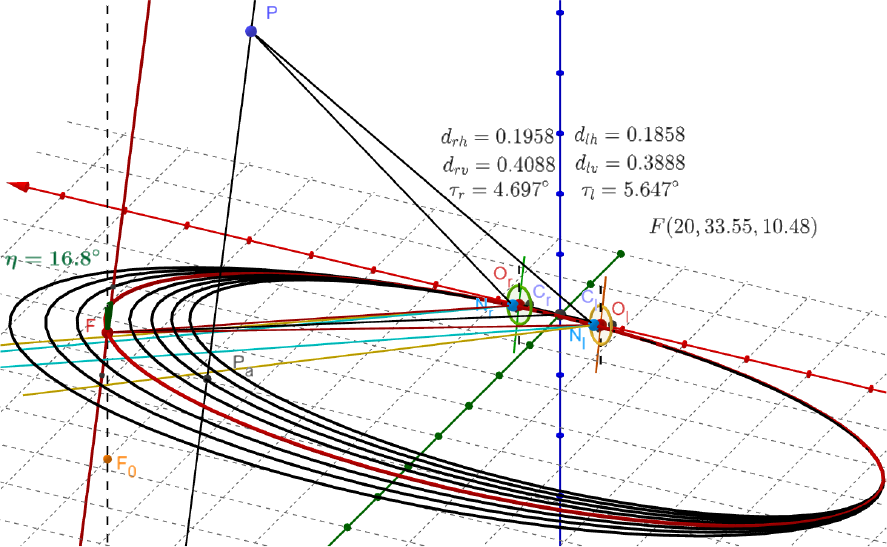
The iso-disparity curves are ellipses for the fixation of the distance less than the abathic distance.

In Figure 6, for the fixation at *F* (*−*20, 158.25, *−*13.48), the horizontal disparity value is *δ*(*P*)_*h*_ = 0.02 cm, the vertical disparity value is *δ*(*P* )_*v*_ = *−*0.002 cm and the torsional disparity value is *δ*_*t*_ = *τ*_*r*_ *−τ*_*l*_ = *−*0.033^*°*^. Also, the subjective vertical horopter is tilted in the plane 1.4*x* + 0.07*y* = *−*17 by *η* = 8.07^*°*^ top-away-relative to the true vertical.

In Figure 7, for the fixation at *F* (20, 33.55, 10.48), the horizontal disparity value is *δ*(*P*)_*h*_ = *d*_*rh*_ *− d*_*lh*_ = 0.01 cm, the vertical disparity value is *δ*(*P* )_*v*_ = *d*_*rv*_ *− d*_*lv*_ = 0.02 cm and the torsional disparity value, *δ*_*t*_ = *τ*_*r*_ *− τ*_*l*_ = *−*0.95^*°*^. Finally, the subjective vertical horopter is tilted bottom-away in the plane 3*x* + 0.05*y* = *−*60 by *η* = 16.8^*°*^.

### 4.4 Disparity Spatial Coordinates Under the Head Roll

In this final subsection, I discuss the special transformation of the disparity lines: starting in the ERP, the head is rotated about the horizontal line passing through the origin of the head coordinates and the point *F*_0_(0, 99.55, 0) such that the fixation point *F*_*a*_(0, 99.55, 3.48) does not change. This is shown in Figure 8.

**Figure 8:**
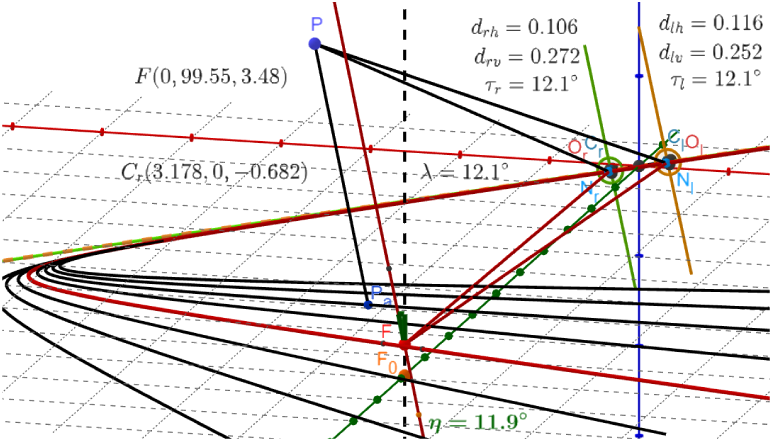
The head is rotated by *λ* = 12.1^*°*^ around the horizontal line passing through the origin of coordinates and *F*_0_(0, 99.55, 0) in the ERP with the fixation point *F*_*a*_ remaining stationary. The disparity straight frontal lines are transformed into hyperbolas, and the subjective vertical horopter is tilted.

In this simulation, the coordinates of the right and left AEs’ centers, *C*_*r*_(−3.25, 0, 0) and *C*_*l*_(−3.25, 0, 0) are rotated by the positive rotation *λ* = 12.1^*°*^. With the fixation point stationary for this head rotational shift, straight frontal lines transform into a family of hyperbolas. The right eye coordinates after rotation are shown in this figure; the left eye rotated coordinates differ in sign. In this case, the horizontal disparity value is *δ*(*P* )_*h*_ = −0.01 cm, the vertical disparity value is *δ*(*P* )_*v*_ = 0.02 cm, and the torsional disparity is zero. The subjective vertical horopter is tilted top-to-right in the plane 0.06*x* + 2*y* = 204 by *η* = 11.9^*°*^.

## 5 Discussion of the Simulated Eyes Postures

The simulations shown in Figures 3 and 4 allow us to formulate the following observations about spatial disparity coordination and the torsional disparity for these cases. The spatial disparity in the ERP is organized by true vertical planes passing through the frontal disparity lines. For any point in the binocular field, the vertical disparity is zero, and the horizontal disparity value is given by the frontal disparity line above which the point is located on the vertical plane. For the true vertical shift of the fixation point from the ERP, the objective binocular space (and vertical planes above disparity lines) tilts by the same amount as the vertical horopter tilts at the fixation point. This visual space is seen subjectively as the same space before the fixation point was shifted. In particular, the vertical and torsional disparities remain zero.

These results indicate that the curvature of visual space vanishes for those eyes postures, extending the 2D results in [Turski, 2023b] for the horizontal visual plane to the 3D results considered here. Moreover, they clearly show that the vertical disparity is not a binocular cue, agreeing with the general evidence provided by the random-dot-stereograms [Julesz, 1971] and Wheatstone stereoscope [Wheatstone, 1838].

The situation is unexpectedly more complicated for the simulating horizontal shifts from the ERP shown in Figure 5. Although the values of the vertical and torsional disparities are small and the hyperbolas are well approximated by straight lines at the fixation, they can affect our impression of being immersed in the ambient environment.

Although eyes shifted horizontally and vertically are in secondary positions, their disparity curves are markedly different after each shift, as it is shown in Figures 4 and 5. My observation is that I use up-and-down eye movement more often without the head movement than when I move my eyes to the left or right. Is this behavioral dissimilarity connected to preserving the straight frontal disparity lines after vertical shifts and transforming them into hyperbolas after horizontal shifts? This question is worth investigating.

The one common feature of the vertical horopter orientations in Figures 5-8 is that under any change either in fixation point or the head position, the vertical horopter is tilted in the different vertical plane in each simulation. This aspect was never considered in experimental observations. The results show that some coefficients in the plane equations are much smaller than the others, which makes planes close to the sagittal or transversal direction for the cases considered in simulations.

In the tertiary posture simulations shown in Figures 6 and 7 and the head roll simulation in Figure 8, the geometry of visual space is much more complicated. Although the curvature of visual space will not vanish, the vertical horopter is still a straight line, but its plane of tilt depends on the fixation point. Although the visual space geometry in the 3D simulations in this study will be difficult to analyze, the results add to other contentions about the relevance of Luneburg’s assertion of the constant hyperbolic curvature visual space [Luneburg, 1947, Blank, 1957].

In the simulation in Figure 8 of the head roll from the ERP when the fixation point does not move, ocular torsion in both eyes is the same and is equal to the head torsion. It does not affect the single vision fusion but may have some effect on balance because the vertical direction of the gravity force differs and does not agree with the subjective vertical in retinal projections of the environment. This simulation of the head roll corresponds to the head tilt test that provokes a compensatory torsional eye movement in the opposite direction to the head motion to keep a more stable image on the retina [Kushner, 2004].

I recall that what distinguishes the simulation of the eye ocular torsion under head roll in the study presented here is the inclusion of the eye’s natural asymmetry of optical components. It should be relevant to experimental investigations.

For example, in [Kushner, 2009], the clinical study of ocular torsion located the precise position of the fovea in the fundus photographs to measure the torsion.

## 6 Constructions of Disparity Conic Sections, Computation of Disparities, and Epipolar Geometry

This section discusses how disparity curves are constructed and how the horizontal, vertical, and torsional disparities are calculated. I also discuss in the binocular system with AEs the epipolar geometry, which simplifies the identification of the two retinal images corresponding to their spatial stimulus.

### 6.1 Disparity Conic Sections and the Vertical Horopter

Figure 9 explains the construction of disparity conic sections for the right eye in the simulating disparity ellipses in Figure 7.

**Figure 9:**
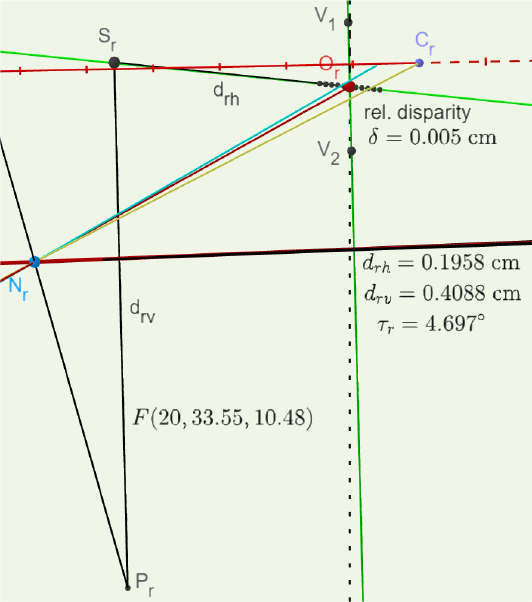
The details of the construction of the disparity ellipses and the vertical horopter in Figure 7 are shown for the right eye. A detailed description is given in the text. The red line starting at *O*_*r*_ and passing through the nodal point *N*_*r*_ is the visual axis, and the orange line from *C*_*r*_ and through *N*_*r*_ is the optical axis. The blue line through *N*_*r*_ is the optical line of the lens, hence, perpendicular to the image plane. The horizontal and vertical coordinates of the projection *P*_*r*_ of the point *P* are *d*_*rh*_ and *d*_*rv*_, respectively. *τ*_*r*_ is the ocular torsion of the right eye.

The horizontal coordinate axis in the image plane for each eye is the intersection of this plane with the visual plane for a given fixation point. The origin of these coordinate axes is the optical center *O*_*r*_ for the right AE and *O*_*l*_ for the left. Following [Turski, 2020], the points on the image planes covering the corresponding retinal points are symmetrically distributed about the optical centers on the horizontal lines for a binocular fixation. They give points of the proximal relative disparity shown in Figure 9 for the right AE spaced around *O*_*r*_ by 0.005 cm. The intersections of the lines through the pairs of corresponding points and the nodal point of each eye give the family of the disparity ellipses, resulting in the distal disparity coordination in the visual plane shown in Figure 7. In this particular case, the subjective vertical horopter is a straight line passing through the intersection of spatial points obtained by the back-projecting pair of points *V*_1_ and *V*_2_ on the vertical green line in the image plane passing through the optical centers *O*_*r*_ or *O*_*l*_ for the right or left AE, respectively. Each pair’s points (*V*_1_ and *V*_2_ shown in Figure 9 for the right eye) are the same distance from the corresponding optical centers.

The vertical horopter exists only for symmetric fixations; otherwise, the intersection of the lines described above in the construction of the vertical horopter is empty. The subjective vertical horopter is extended to a central range of azimuthal angles by using the three-dimensionality of this horopter, first emphasized by Amigo in [Amigo, 1974]. He studied the stereoscopic sensitivities of the retinal regions at the various elevations above and below the fovea, which corresponds to Panum’s fusional region.

The vertical horopter’s extension is done as follows. The two back-projected rays used for constructing the vertical horopter in the resting eyes posture, each in a different eye, are replaced with cylinders. The other two rays intersect the cylinders, and the vertical horopter passes through the midpoints of the intersections. The cylinder’s radius determines the vertical horopter range of the azimuthal angles. In the simulations, I assumed a 1 cm radius at the abathic distance of about 100 cm and 10 cm above and below the fixation point.

### 6.2 Torsional Disparity

The geometric construction in *GeoGebra’s* simulation shown in Figure 10 computes the two eyes’ ocular torsions for the case in Figure 7 with the fixation point denoted here by *F* ^1^. By choosing the appropriate inputs of the fixation point, the ocular torsions were calculated for all simulations shown in Figures 3-8.

**Figure 10:**
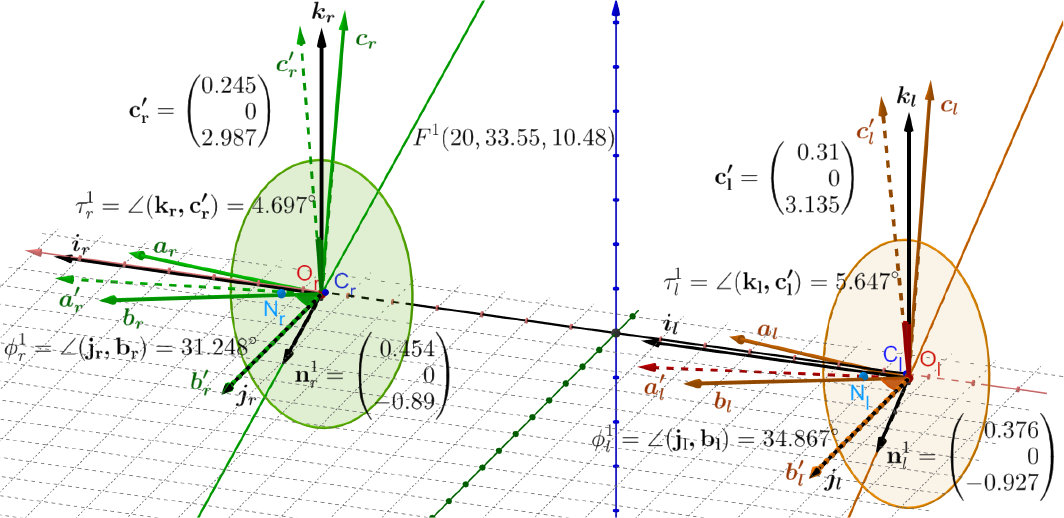
The black frames in the ERP (i_r_, j_r_, k_r_) and (**i**_**l**_, **j**_**l**_, **k**_**l**_) for the right and left AEs are attached at the image planes optical centers *O*_*r*_ and *O*_*l*_. Their orientations agree with the upright head’s frame. The solid-colored frames are moving from the position of black frames for a given fixation *F* ^1^. The dashed-colored frames are obtained by rotating the solid-colored frames by moving **b**s vectors onto the **j**s vectors. The rotation axes are given by 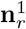 and 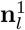, with theangle of rotations 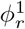 and 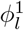, are contained in the co-planar image planes of the ERP, parallel to the head frontal plane.The ocular torsions of the AEs are 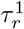 and 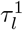.

In Figure 10, the black frames (**i**_**r**_, **j**_**r**_, **k**_**r**_) and (**i**_**l**_, **j**_**l**_, **k**_**l**_) for the right and left eyes are attached at the image planes’ optical centers *O*_*r*_ and *O*_*l*_ and agree with the upright head’s frame in the ERP. The solid green frame for the right AE, (**a**_**r**_, **b**_**r**_, **c**_**r**_), and the brown frame for the left AE, (**a**_**l**_, **b**_**l**_, **c**_**l**_), are moving with the image planes from their position that initially agrees with the respective black frame for each AE.

To calculate each eye’s ocular torsion, the green and brown frames are rotated such that **b**s vectors are overlaid with **j**s vectors. The rotations are in the planes spanned by **b**s and **j**s vectors such that rotation axes 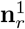 and 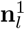 are perpendicular to these planes. The rotations result in two “primed” frames shown in dashed lines. The rotations angles, ∠(**j**_**r**_, **b**_**r**_) = 31.248^*°*^ and ∠(**j**_**l**_, **b**_**l**_) = 34.867^*°*^ are denoted by 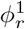and 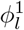. Because each frame’s rotation axes 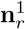 and 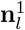 are perpendicular to **j**_**r**_ and **j**_**l**_, which are parallel to the head’s medial axis, they are in the coplanar image planes for the ERP. The ocular torsion, defined by the angles 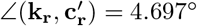 and 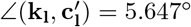, are denoted by 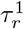 and 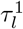 such that the torsional disparity 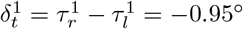. All counterclockwise right-hand rotations are positive.

The results presented here establish the geometrical calculations of the eyes’ ocular torsions for any binocular fixation reached from the ERP. I stress that the ERP defined only for bifoveal fixations provides a neurophysiologically motivated definition for Listing’s law eye’s primary position that has been originally intended for the single eye [Turski, 2020].

The relation between the ocular torsion calculated here and Listing’s law which is commonly used in the control of 3D eye orientations, will be discussed in Section 7.

### 6.3 Epipolar Geometry for AEs

The epipolar geometry is constructed in the image planes of the AEs in Figure 11.

**Figure 11:**
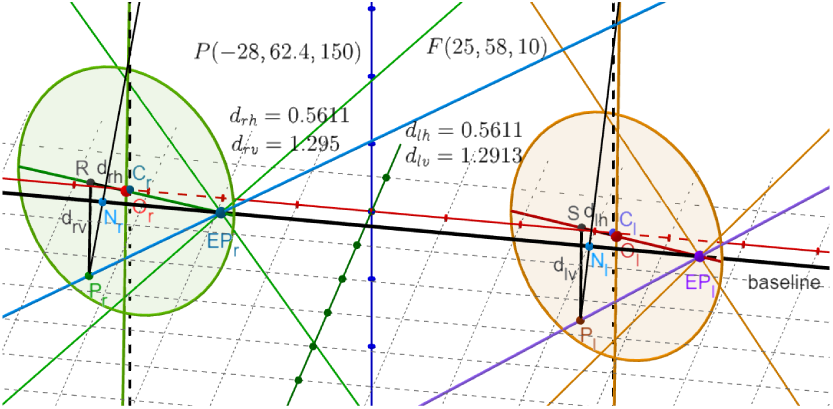
The projections of the point *P* to the respective image planes, *P*_*r*_ and *P*_*l*_, have the coordinates *d*_*rh*_ and *d*_*rv*_ for the right eye and *d*_*lh*_ and *d*_*lv*_ for the left image plane. The black line through the nodal points *N*_*r*_ and *N*_*l*_ is the baseline of epipolar geometry. The epipolar lines: two green and blue lines intersect at the epipole *EP*_*r*_, and two brown and purple lines intersect at the epipole *EP*_*l*_. The blue line passes through *P*_*r*_, and the purple line passes through *P*_*l*_.The epipolar geometry for the binocular system with AEs is discussed in the text.

Each circle represents the projection through the nodal point of the binocularly useful part of the retina into the respective image planes. Please refer to [Hartley and Zisserman, 2003] for an introduction to epipolar geometry in computer vision, as it is closely related to the presentation in the binocular system with AEs.

In Figure 11, the black line passing through the centers of projections—the nodal points *N*_*r*_ and *N*_*l*_—is the baseline in the epipolar geometry. The epipolar lines are constructed in the ERP by the intersection of the planes containing the baseline with the image plane of each AE. When the fixation point moves from *F*_*a*_ to *F*, the nodal points move, and the image plane rotates accordingly. The epipolar lines in the image planes, under this change of this eyes’ fixation, intersect at the epipoles on the baseline. In Figure 11, for the fixation *F* (25, 58, 10), three epipolar lines in each image plane, two green and one blue in the right image plane and two brown and one purple in the left image plane, intersect at epipoles *EP*_*r*_ and *EP*_*l*_ on the baseline. The blue line contains *P*_*r*_, and the purple line contains *P*_*l*_, the projections of the spatial point *P* into the right and left image planes.

## 7 Discussion of the Relation to Listing’s Law

Historically, the torsion of the eye was determined by Donders, Listing, and Helmholtz between 1848 and 1867. Donders’s law is a general statement that with the head erect and looking at infinity, any gaze direction has a unique torsional angle, regardless of the path followed by the eye to get there. Listing’s law (L1), formulated by Helmholtz in 1867, restricts Donders’s law by stating that visual directions of sight are related to rotations of the eye so that from the reference position, all rotation axes lie in a plane, today referred to as the displacement plane [Tweed and Vilis, 1990]. When the reference orientation is perpendicular to the displacement plane, the plane is referred to as Listing’s plane, and the unique reference direction is the primary direction. However, the displacement plane’s orientation was never precisely specified, as is discussed below.

The Listing’s law has been extended in two ways. First, Listing’s plane must tilt by half the angle of eye eccentricity from the primary direction [Mok et al., 1992, Minken and van Gisbergen, 1994]. It is known as the “half-angle” rule. Second, for the eyes’ posture with converging gazes on a near fixation point, each eye’s displacement plane of the L1 law turns by the amount proportional to the gaze direction angle [Van Rijn and Van der Berg, 1993, Tweed, 1997]. It is known as binocular Listing’s law (L2).

The problem with Listing’s laws is the lack of clear separation between the theorems of ocular rotations and the constraints imposed by the ocular plant and neural processing. This follows from an original formulation of Listing’s law for the single eye, with extension to “binocular” conjugate eye movements with parallel lines of sight of both eyes. Then, the binocular L2 law is formulated by the Listing planes tilted by angles proportional to the initial vergence angle. However, values of the proportionality coefficients are controversial [Van Rijn and Van der Berg, 1993]. Further, the primary position, which is still a part of L2 law, corresponds to the eyes’ movements with parallel gazes, and, consequently, its neurophysiological significance has remained elusive despite its theoretically important underpinning of the oculomotor research [Hess and Thomassen, 2014].

How do Listing’s law and its extensions compare to my study?

The geometric theory developed here for binocular fixations is fundamentally based on Euler’s rotation theorem. This theorem states that any two orientations of a rigid body with one of its points fixed differ by a rotation about an axis specified by a unit vector passing through the fixed point. I recall that fixed-axis rotations are lines of the shortest length (geodesics) on the Lie group *SO*(3) of rotations and, hence, are *optimal*, for example, [Novelia and O’Reilly, 2015]. Here, Euler’s rotation theorem is used in the framework of Rodrigues’ vector to each eye constrained by the eyes’ bifoveal fixations.

To introduce this framework, I start with the conclusion from Euler’s rotation theorem: Any rotation matrix *R* can be parametrized as *R*(*ϕ*, **n**) for a rotation angle *ϕ* around the axis **n**. This parametrization is unique if the orientation of 0 *< ϕ <* 180^*°*^ is fixed. Usually, a counterclockwise right-hand orientation is chosen to get angles’ positive values.

The pair *ρ* = cos(*ϕ/*2), **e** = sin(*ϕ/*2)**n** is known as *Euler-Rodrigues parameters* and

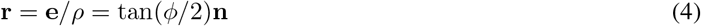

as *Rodrigues’ vector* usually referred to as *rotation vector* [Piña, 2011]. Rodrigues proved that under composition *R* = *R*_2_*R*_1_, the corresponding Euler-Rodrigues parameters transform as follows [Gray, 1980],

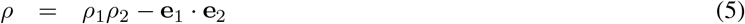

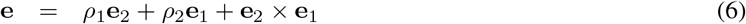

Then, it is easy to see that the corresponding composition for Rodrigues’ vectors is

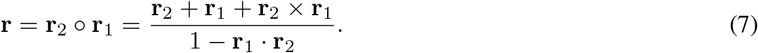

Further, **r**^*−*1^ = *−***r** and tan(*−ϕ/*2)**n** = tan(*ϕ/*2)(*−***n**) define the same rotation vector.

Note that (cos(*ϕ/*2), sin(*ϕ/*2)**n**) is a unit quaternion that describes the rotation by *ϕ* around **n**, and Rodrigues used the quaternions to obtain (5) and (6) because in his time vectors were still not invented.

The rotation vectors in the simulation shown in Figure 10 for fixation *F* ^1^(20, 33.55, 10.48) are

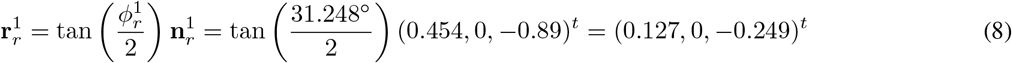

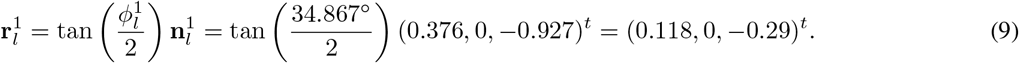

for the right and left eye, respectively.

Now, I can compute directly the torsion changes between tertiary eyes’ binocular fixations, which usually are obtained in oculomotor research using Listing’s L2.

To this end, I first show in Figure 12 the simulation for the change of the fixation points from the resting eyes’ posture at *F*_*a*_(0, 99.55, 3.48) to *F* ^2^(15, 53.55, 20.48). Following similar calculations for *F* ^2^ as before for the fixation *F* ^1^, from results presented in Figure 12, I obtain,

**Figure 12:**
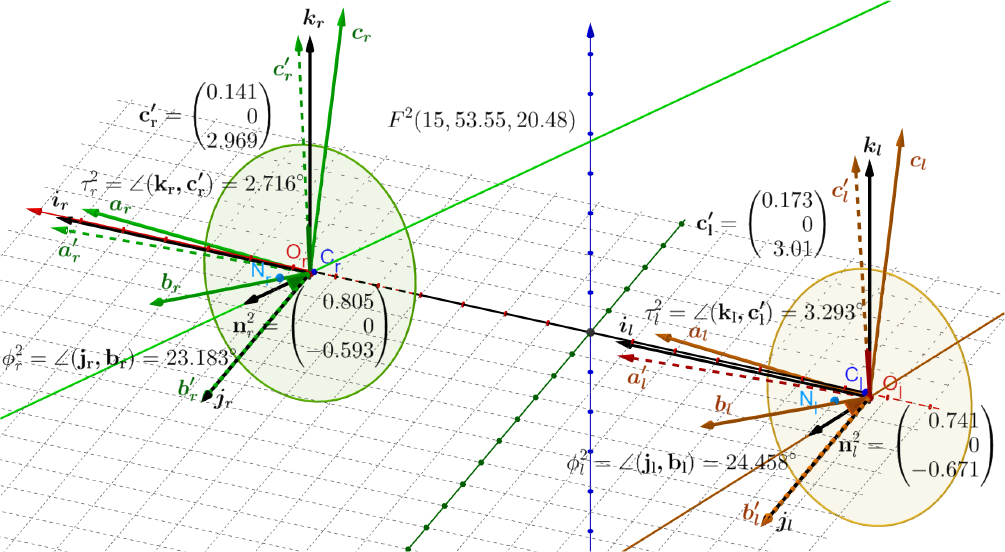
For the fixation *F* ^2^(15, 53.55, 20.48, the frames’ angles of rotations around the axes 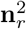 and 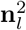 are 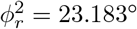 and 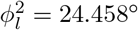, respectively for the right and left AE.

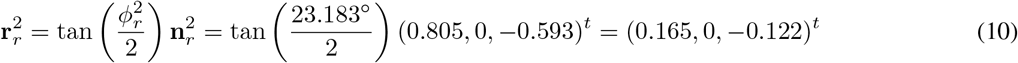

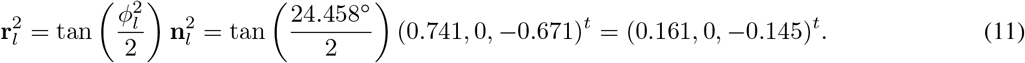

Then, frames rotations that align vector **b**_*r*_ for *F* ^1^(20, 99.55, 10.48) with with vector **b**_*r*_ for *F* ^2^(15, 53.55, 20.48) and similarly for vectors **b**_*l*_s are calculated as follows. Let these Rodrigues’ vectors be denoted by 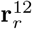 and 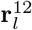 for the right and left eye, respectively. Thus, using properties of Rodrigues’ vectors discussed above, I obtain

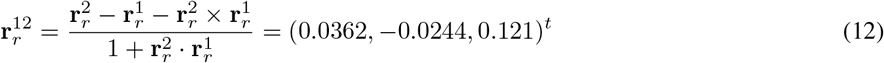

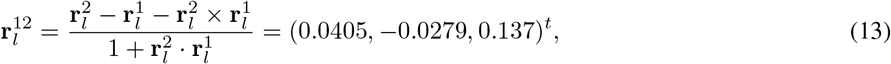

so that from (4), (12) and (13), I have

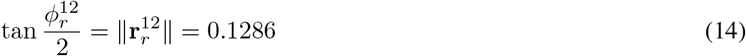

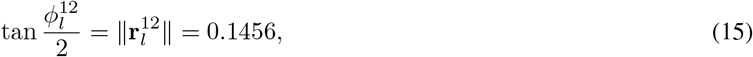

which gives the corresponding angle values 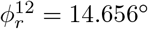 and 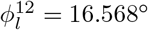 for the right and left eye, respectively

Finally, these Rodrigues’ vectors can be written as

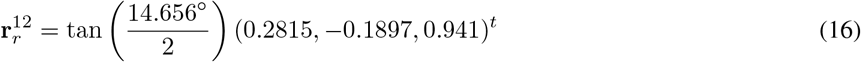

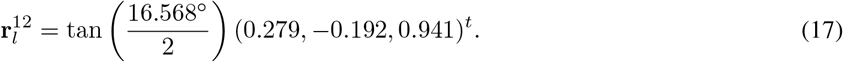

and the ocular torsions when the eyes fixation changes from *F* ^1^ to *F* ^2^ can be calculated as 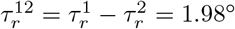and 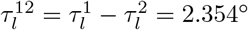 using the values given in Figures 10 and 12.

I conclude this section with the “half-angle” rule usually attributed to the non-commutativity of 3D rotations, see [Martinez-Trujillo, 2005], for example. The Listing plane, which can only be precisely discussed for the two eyes’ position with parallel gaze directions, is represented in the binocular system with the AEs by the ERP. I recall that the AEs’ image planes for the resting eyes are coplanar and parallel to the transverse head plane for the bifoveal fixation *F*_*a*_(0, 99.55, 3.48) corresponding to the average abathic distance fixation measured for empirical horopters. When the eyes fixation changes from *F* ^1^(20, 33.55, 10.48) to *F* ^2^(15, 53.55, 20.48), the half-angle rule is demonstrated in Figure 13 by using Rodrigues’ vectors (16) and (17).

**Figure 13:**
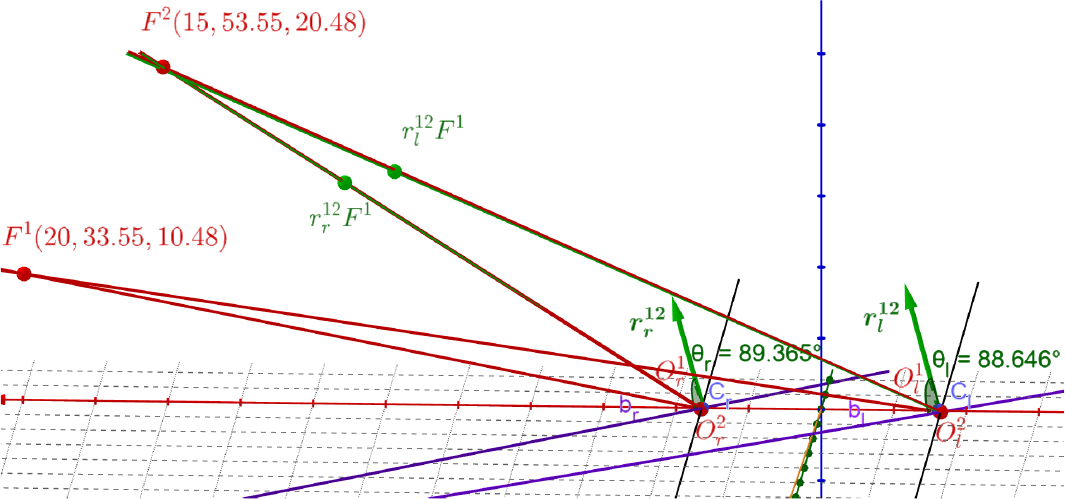
The half-angle rule for the fixation changing from *F* ^1^ to *F* ^2^. Rodrigues’ vectors 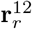 and 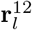 rotate visual axes for *F* ^1^ to align them with the visual axes for *F* ^2^. Green dots 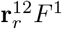 and 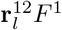 are rotations of *F* ^1^ by the indicated Rodrigues’ vectors. The bisectors of the angles between the visual axes of *F* ^1^ and the optical axes of eyes are denoted as purple lines *b*_*r*_ and *b*_*l*_ for the right and left eye, respectively. The angles between the bisectors and Rodrigues’ vectors, *θ*_*r*_ and *θ*_*l*_, are close to 90 degrees. The small discrepancy is due to the eye’s asymmetry parameters.

Clearly, the “half-angle” rule follows from Euler’s rotation theorem here implemented in the framework based on Rodrigues’ vectors. The small discrepancy discussed in the caption of Figure 13 is the result of the eye’s asymmetry of optical components when the rotations are applied to visual axes rather than as originally defined for alignment of **b**_*r*_s vectors and alignment of **b**_*l*_s for *F* ^1^ and *F* ^2^ fixations. Importantly, the nonzero vergence coefficients that define the tilts of the initial “Listing plane” for each eye in the binocular extensions L2 that are controversial are not needed in the framework used in this study.

## 8 General Discussion

The recent studies in [Turski, 2018, Turski, 2020, Turski, 2023b] established in *GeoGebra*’s dynamic geometry environment the first geometric theory of longitudinal iso-disparity curves supported by the eye’s anatomy and physiology. The study presented here extended this theory to a 3D framework and critically assessed its impact on Listing’s law, which is important in the control of 3D eye movements and is used in clinical implications for strabismus and the optimal management of this disorder [Wong, 2004]. In this section, I will briefly describe the main contributions of the above-mentioned papers.

One of the main contributions of [Turski, 2018] was the realization that the asymmetry of retinal correspondence results from the healthy human eyes’ asymmetry of optical components: the fovea is displaced from the eyeball’s posterior pole, and the lens is tilted relative to the cornea [Chang et al., 2007, Schaeffel, 2008]. The functional significance of this asymmetry is that the lens tilt partly compensates for the optical aberrations resulting from the fovea’s anatomical displacement on the retina from the posterior pole and the asphericity of the cornea [Artal, 2014]. The AE model, introduced in [Turski, 2018], comprises foveal displacement and lens tilt angles.

It was demonstrated in [Turski, 2020] that the projection of the asymmetric distribution of corresponding retinal elements into the image planes of the AEs in the ERP is symmetric. Thus, the unknown asymmetry of the retinal correspondence is fully specified in the image plane in terms of the AE parameters specifying the human eye asymmetry of the optical components. This property establishes the physiological origin of retinal correspondence asymmetry for the first time.

In [Turski, 2023b], the binocular system with AEs was extended to the iso-disparity conic sections in the horizontal visual plane formulated in terms of the retinal relative disparity formulated on the AE’s image plane. These disparity curves provide spatial coordination of the retinal disparity, which is fundamental in understanding the perceived objects’ spatial relations and shape, i.e., visual space geometry and stereopsis. The lack of disparity spatial coordinates that would more richly describe the structure of binocular visual space was mentioned in [Arditi, 1986]. In the present study, the horizontal, vertical, and torsional disparities are similarly calculated in the binocular system with the AE model augmented by the vertical asymmetry of optical components in addition to previously considered horizontal asymmetry.

Ocular torsion has been a controversial issue in contemporary vision science, as exemplified by the use of “true torsion” and “false torsion.” By usual definitions, true torsion, or torsion, is the eyeball’s rotation about the gaze direction, while false torsion mainly results from the dependence of perspective projection on gaze orientation. In ophthalmology, the false torsion is described as the discrepancy between the vertical corneal meridian and the objective vertical [Noorden von and Campos, 2002]. Describing ocular movement either in Helmholtz or Fick coordinates results in different values of false and true torsions for the same eye orientation.

However, false torsion vanishes in the framework of quaternions or Rodrigues’ (rotation) vectors. The framework of Rodrigues’ vector is used in this study. The present study contributes to the unambiguous understanding of the eye’s ocular torsional movement based on purely geometrical consideration but in the anatomically supported binocular system with AEs. Referring to [Turski, 2020], I recall that the ERP used in the binocular system with AEs corresponds numerically to the abathic distance fixation measured for empirical horopters. Further, both are within the range of the eye muscles’ natural tonus resting position distance [Jaschinski-Kruza, 1991], which serves as a zero-reference level for convergence effort [Ebenholtz, 2001].

Thus, the ERP provides a compelling alternative for the imprecise definition of the primary eyes’ position, an integral part of Listing’s law constraining the eye’s torsional orientation. Originally intended for a single eye, this primary position is defined as both eyes being directed straight ahead by an upright head. Then, in the theory of binocular oculomotor research, the eye movement is constrained to conjugate eye movements with parallel lines of sight of both eyes, also referred to as “double eye” movement [Hepp, 1990]. The definition of primary position is the reason why its neurophysiological significance remains elusive despite its theoretical importance to oculomotor research [Hess and Thomassen, 2014].

In the geometric theory presented here and supported by simulations, the consecutive change in the eyes’ binocular posture for the fixed upright head is precisely described as (1) a change relative to the binocular ERP or as (2) shifts between binocular tertiary postures. Both avoid the ambiguous “double eye” movement with parallel gaze directions.

The first description of the consecutive eyes’ binocular posture shifts relative to the ERP, which should be more economically executed by the vision system, may be supported by experimental observations. In [Jampel and Shi, 1992], the experimental measurements supported by photographic and video analysis demonstrated that the anatomically determined primary position is a natural constant position to which the eyes are automatically reset from any displacement of the visual line. Further, the evidence was presented to indicate an active neurologic basis for the primary position.

In the second description, the eye’s binocular posture changes between two tertiary fixations. It verified the half-angle rule, which has Euler’s rotation theorem origin and can be formulated as a theorem with a small discrepancy attributed to the asymmetry of the eye’s optical components. Moreover, the coefficients of proportionality between the vergence and the tilts of the primary eye positions used in Listing’s L2 law that are controversial are not needed.

How do I explain the differences between an unambiguous formulation in my work and a controversial formulation of binocular Listing’s law? Ruete first mentioned in 1855 what we now know as Listing’s law in his textbook on ophthalmology and attributed it to Johannes Benedict Listing. This law was formulated for the single eye. Its significance was established after Helmholtz’s measurements of visual afterimages at various eye positions verified it. Today, Listing’s law is called L1, and its binocular extension is L2. However, as discussed before, L2 is highly controversial. Moreover, the precise consideration of the L2 law had to be restricted to the above-mentioned “double eye” movements. In contrast, my approach is binocular from the beginning and precise by employing Euler’s rotation theorem in Rodrigues’ vector framework, which ensures the *optimality* of the eyes’ binocular movement. As a German mathematician, Listing could base this law on Euler’s rotation theorem, which was lost during the communication between a mathematician and an ophthalmologist.

The study presented here provides for the first time the comprehensive construction of retinal disparity’s spatial coordinates for eyes’ secondary and tertiary binocular postures, and the head rolled around its medial axis with the stationary point of fixation. The resulting coordinates allow computations of longitudinal and vertical disparities of spatial visual stimuli. Further, torsional disparity for eyes’ postures is computed in the framework of Euler’s rotation theorem. It lays the groundwork for studying visual space geometry and assessing the geometric and ocular motor plant or neural constraints to Listing’s law.

## Supporting information

3D applet

## Notes

### Competing Interest Statement

The authors have declared no competing interest.

### Summary of Updates

a small correction of the statement about geodesics on page 11

