## Supplementary material for "A Critical Study in Stereopsis and Listing’s Law": 3D applet: 3D applet.pdf

**For**

*Jacek Turski*

### Applet: Iso-disparity Conics Transformation with Eyes Posture in *GeoGebra*

To run the applet, you need install the GeoGebra Classic 6. It is free and easy to do. Then, follow the link: https://www.geogebra.org/m/yh4fvg34

The initial screen in the applet shows the resting eyes' posture is shown in Fig. 1. For visualization purposes, in this applet, the dimensions are not anthropomorphic as they are in the article.

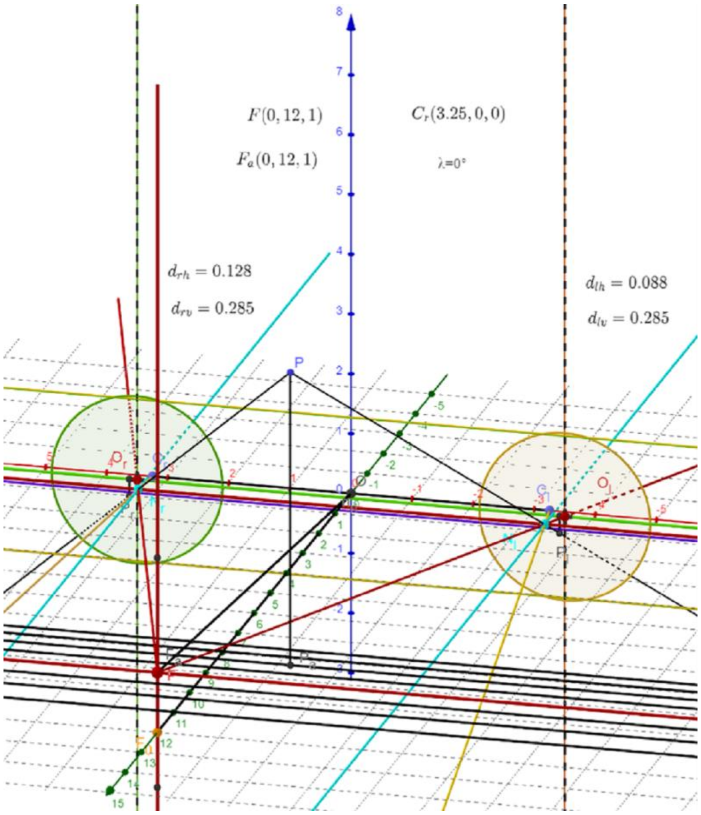


Fig. 1. The initial applet's screen.

The minimal relative disparity in AE of 0.02 cm is back-projected to the space giving the same disparity between consecutive iso-disparity spatial lines. The initial horizontal disparity of the spatial point $P$ over the second iso-disparity line from the horopter is $d_{rh} - d_{lh} = 0.128 - 0.088 = 2(0.02)$ cm and the vertical disparity is zero. The torsional disparity is not included in this simulation.

To simulate the iso-disparity transformations, highlight in the applet the red dot $F$ (fixation). You will be able to move it horizontally or vertically. As you move it, the initial iso-disparity frontal lines in the eyes' resting posture and the subjective vertical horopter deform accordingly, and the AE horizontal and vertical coordinates of projections of P are displayed. To reset, return the red dot to the (smaller) blue dot $F_a(0,12,1)$, the resting eyes' posture fixation. You will see that the text window $F(a,b,c)$ will reset to $F_a$.

To rotate the head about the viewing direction in the resting eyes' posture, highlight the blue dot $C_r(3.25,0,0)$ and move it. You will see the deformed iso-disparity conics and in the text window the coordinates of $C_r$ and the angle of the head's tilt $\lambda$. Move the $C_r$ point back to the original position.

You should not make more than two consecutive movements of the fixation point because you will face the problem of returning to the starting position as you view only 2D projections of the 3D scene. You should rather move to any location and investigate 3D geometry by rotating the view before returning to the initial resting eyes' posture. Then move the (red) fixation $F$ to another location.

The black line through the nodal points $N_r$ and $N_l$ is the baseline in the epipolar geometry. The green lines at the top and bottom of the green circle, the brown lines at the top and bottom of the brown circle, blue line through $P_r$, and purple lime through $P_l$ are the epipolar lines. While running the applet, for fixation points outside of the midsagittal plane, all the epipolar lines intersect at the epipoles $EP_r$ and $EP_l$ located on the epipolar baseline.
